# Speak and you shall predict: speech at initial cocaine abstinence as a biomarker of long-term drug use behavior

**DOI:** 10.1101/2023.07.18.549548

**Authors:** Carla Agurto, Guillermo Cecchi, Sarah King, Elif K. Eyigoz, Muhammad A. Parvaz, Nelly Alia-Klein, Rita Z. Goldstein

**Affiliations:** IBM Research, 1101 Kitchawan Rd, Yorktown Heights, NY, 10598; Psychiatry and Neuroscience Departments, Icahn School of Medicine at Mount Sinai, 1 Gustave L. Levy Place, New York City, NY, 10029; Artificial Intelligence and Human Health, Icahn School of Medicine at Mount Sinai, New York City, NY, 10029

## Abstract

**Importance:** Valid biomarkers that can predict longitudinal clinical outcomes at low cost are a holy grail in psychiatric research, promising to ultimately be used to optimize and tailor intervention and prevention efforts.

**Objective:** To determine if baseline linguistic markers in natural speech, as compared to non-speech clinical and demographic measures, can predict drug use severity measures at future sessions in initially abstinent individuals with cocaine use disorder (iCUD).

**Design:** A longitudinal cohort study (August 2017 – March 2020), where baseline measures were used to predict outcomes collected at three-month intervals for up to one year of follow-up.

**Participants:** Eighty-eight initially abstinent iCUD were studied at baseline; 57 (46 male, age 50.7+/-7.9 years) came back for at least another session.

**Main Outcomes and Measures:** Outcomes were self-reported symptoms of withdrawal, craving, abstinence duration and frequency of cocaine use in the past 90 days at each study session. The predictors were derived from 5-min recordings of vocal descriptions of the positive consequences of abstinence and the negative consequences of using cocaine; the baseline cocaine and other common drug use measures, demographic and neuropsychological variables were used for comparison.

**Results:** Models using the non-speech variables showed the best predictive performance at three(r>0.45, *P*<2×10^-3^) and six months follow-up (r>0.37, *P*<3×10^-2^). At 12 months, the natural language processing-based model showed significant correlations with withdrawal (r=0.43, *P*=3×10^-2^), craving (r=0.72, *P*=5×10^-5^), days of abstinence (r=0.76, *P*=1×10^-5^), and cocaine use in the past 90 days (r=0.61, *P*=2×10^-3^), significantly outperforming the other models for abstinence prediction.

**Conclusions and Relevance:** At short time intervals, maximal predictive power was obtained with models that used baseline drug use (in addition to demographic and neuropsychological) measures, potentially reflecting a slow rate of change in these measures, which could be estimated by linear functions. In contrast, short speech samples predicted longer-term changes in drug use, implying deeper penetrance by potentially capturing non-linear dynamics over longer intervals. Results suggest that, compared to the common outcome measures used in clinical trials, speech-based measures could be leveraged as better predictors of longitudinal drug use outcomes in initially abstinent iCUD, as potentially generalizable to other substance use disorders and related comorbidity.

**Key Points:** *Question:* Can natural language processing be leveraged to predict longitudinal drug use outcomes in individuals with substance use disorder?

*Findings:* In this prospective study that included initially abstinent individuals with cocaine use disorder (iCUD), models using demographics, neuropsychological measures and drug use patterns at baseline were compared to those using minimally structured short natural speech samples relating the positive consequences of abstinence and the negative consequences of using drugs, showing a differential prediction of outcomes measured up to one year later. At three and six months, the former outperformed speech models, including approximately 50% of the variability in craving and 40% in abstinence duration. At 12 months from baseline, speech models were superior, predicting 50% of the variability in abstinence duration.

*Meaning:* Speech variables derived through natural language processing can predict clinically meaningful drug use outcome measures in addiction, with greater value at longer intervals. The applicability of language modeling to aid in assessing treatment response and risk in drug addiction warrants further investigation in clinical settings.

## Introduction

Identifying predictive markers of short and long-term outcomes in psychiatric populations is of foremost importance for treatment planning and prevention. In particular, accurate longitudinal outcome prediction can effectively inform resource distribution and treatment strategies, enriching and optimizing clinical trials. Maximizing the predictive efficacy of clinically informative measures necessitates dense sampling in diverse populations ^1^. Recruiting numerous cognitive and emotional neurobiological processes, language represents a dense and ubiquitous, yet underutilized, resource for accessing unique markers with potential predictive value for numerous clinical outcomes in human psychopathology. In substance use disorders (SUD), interventions to guide recovery commonly employ language-based strategies (e.g., recounting one’s own story, providing concrete guidance to others, in group settings such as Alcoholics Anonymous/Narcotics Anonymous meetings); lab-developed emotional regulation techniques (e.g., with reappraisal of drug cues or savoring of alternative reinforcing cues) similarly rely on the story people tell themselves when processing a salient cue ^2, 3^. Spoken and written language is also central to clinical evaluations of SUD, which intend to ascertain, using self-report, the amount, frequency, and severity of drug use, encompassing symptoms such as craving and withdrawal, and more general attitudes towards one’s well-being and quality of life. Moreover, drug-biased language has been shown to provide a window into brain function in people with SUD ^4^ potentially serving as a proxy of immediate and life-long learning, a readily accessible behavioral marker of neural plasticity. Therefore, applications of natural language processing (NLP) could hold a considerable potential for enhancing the arsenal of objective behavioral markers for outcome prediction in clinical trials for SUD ^5^.

In SUD research and considering the ongoing opioid crisis, unsurprisingly NLP methods have most commonly used electronic health records to detect and predict problematic substance use, dependence, and treatment outcomes including relapse, in both adult ^6–9^ and pediatric populations ^10^. In cocaine use, NLP was similarly applied to medical/health records and/or clinicians’ notes to identify SUD and guide treatment ^11–14^ and to accurately classify overdose deaths ^15^. Another relevant application is the use of social media data (e.g., Facebook, Reddit) to predict SUD ^16^ and characterize language features (e.g., emotional word use) unique to substance use cessation and abstinence ^17^. More closely aligned to the goals of the current study, NLP-based topic modeling of healthy participants’ written narratives of their experience (quitting or reducing use of recreational drugs following use of a psychedelic drug) predicted long-term substance consumption with approximately 65% accuracy ^18^. Here for the first time, we explore the use of similar NLP techniques of natural speech to longitudinally predict drug use outcomes in individuals diagnosed with SUD.

Clinical trials in SUD typically consider the complete absence, or a continual reduction, of drug use as their primary outcome measures. However, the early stages of recovery are often characterized by nonlinear/dynamic patterns of drug use and cue reactivity, the latter characterized by initial increases followed by decreases as a function of time into abstinence ^19^. In identifying effective outcome predictors, it is therefore important to consider the potential distinctions in the trajectories of short and long-term recovery. Psychometric and behavioral measures including goal-directed thinking ^20, 21^, impulsivity ^22^, and behavioral flexibility ^23^ have previously demonstrated predictive efficacy in individuals receiving treatment for SUD; compared to traditional clinical and demographic variables, these cognitive measures are better predictors of future drug use and treatment retention at both shorter (<six months) ^24–26^ and longer (≥six months) ^27, 28^ intervals, supporting further investigation of such brain function measures. Because natural speech is both scalable and amenable to data collection outside the laboratory (e.g., in clinical and home settings), NLPs using these data may offer specific advantages over other laboratory-based behavioral measures in SUD outcome prediction.

We previously applied NLP and acoustic analysis of natural speech to detect concurrent drug use in healthy individuals and iCUD ^29, 30^. Here for the first time we used BERT ^31^, a deep learning model for language processing, previously applied primarily to detect SUD from social media ^32^, to analyze short speech samples recorded from well-phenotyped initially abstinent iCUD to predict withdrawal, craving, abstinence duration, and 90-day cocaine use at 3, 6, 9, and 12 months after baseline. Specifically, using BERT sentence embeddings, we computed the similarity between the speech samples and inventories (selected based on clinical knowledge) probing individuals’ severity of dependence, cocaine craving, quality of life, and current emotional state. These similarity scores were used in regression models for outcome prediction, evaluated alongside separate models that contained baseline clinical (including the same to-be-predicted drug use measures), demographic, and neuropsychological information (i.e., non-NLP measures commonly used for prediction purposes), for direct comparisons.

## Methods

### Participants

This study was approved by the Institutional Review Board of the Icahn School of Medicine at Mount Sinai, and all participants provided written informed consent. A total of 88 iCUD, initially abstinent as verified by urine testing at the baseline session, were recruited (baseline abstinence durations ranged from 4 to 242 days) using posted flyers, newspaper and other (e.g., Craigslist) ads (including those posted around SUD treatment/rehabilitation facilities), and by word of mouth. Participants came back every three months for one year. In all sessions, measures were acquired for the four dependent outcome variables of interest: 1) withdrawal symptoms in the past 24 hours using the Cocaine Selective Severity Assessment (CSSA) ^33^; 2) craving symptoms with the 5-item Cocaine Craving Questionnaire (CCQ) (Tiffany et al., 1993); 3) duration of current abstinence in days; and 4) days of cocaine use in the past 90 days using the Timeline Followback interview ^34^. At each session, we also recorded participants’ speech about the positive consequences (PC) of abstinence and the negative consequences (NC) of using cocaine, 5 minutes each. See supplemental material for further details on the clinical diagnostic interviews/tools and the speech samples (including verbatim instructions). Participants who completed two or more sessions were included in the analysis (N=57: 46 male, 11 female). Descriptive statistics for all variables collected at baseline are presented in Table 1.

**Table 1.** Demographics, neuropsychological and drug use variables assessed at the baseline visit.

|  | Category | Values<sup>e</sup> |
| --- | --- | --- |
| **Demographics** | Sex | 46 males/11 females |
|  | Age (years) | 50.66 +/- 7.86 |
|  | Education (years) | 12.18 +/- 2.59 |
|  | Race | White: 4, Black: 48, Other: 5 |
| **Neuropsychology** | Verbal IQ | 95.75 +/- 11.92 |
|  | Non-verbal IQ | 9.84 +/- 3.19 |
|  | Depression (BDI<sup>a</sup>) | 9.4 +/- 7.99 |
| **Cocaine, alcohol and cigarette use** | Cigarette smokers | Never: 7, past: 8, current: 32, unknown: 10 |
|  | SMAST<sup>b</sup> (alcohol use) | 4.43 +/- 2.33 |
|  | Cocaine administration | 12 Oral, 44 smoking, 1 injection |
|  | Cocaine age of onset (years) | 23.4 +/- 7.7 |
|  | Cocaine regular use (years) | 17.9 +/- 9.4 |
|  | SDS<sup>c</sup> Dependence severity | 10.42 +/- 3.55 |
| **Cocaine use metrics at baseline (dependent variables)** | Withdrawal (CSSA<sup>d</sup>) | 18.75 +/- 12.97 |
|  | Cocaine craving | 14.34 +/- 11.3 |
|  | Days of abstinence | 48.4 +/- 49.42 |
|  | Cocaine 90 days | 13.18 +/- 15.82 |

<sup>a</sup>BDI refers to Beck's depression index,
<sup>b</sup>SMAST to Short Michigan Alcohol Screening Test,
<sup>c</sup>SDS to Severity of Dependence Scale,
<sup>d</sup>CSSA to Cocaine Selective Severity Assessment
<sup>e</sup>Values in the table are represented as means +/- standard deviation values.

### Analytical Approach

Recordings were automatically transcribed using a customized version of the Speech to text IBM API ^35^. Sentences were identified using NLTK sentence tokenizer and were embedded using RoBERTa ^31^. Similarity scores were computed between these sentences and sentences from three different inventories as shown in Figure 1. Specifically, we selected three sets of statements for which we also computed sentence embeddings: 1) “cocaine-related” set included all five items from the Severity of Dependence Scale (SDS) ^36, 37^, all 10 items from the CCQ-brief (^38^, different than the one administered during the sessions), and eight items a priori determined to assess drug craving under different emotional contexts (e.g., “’I want to use cocaine when I feel anxious”); 2) “emotion-related” set included the 40 items of the State-Trait Anxiety Inventory (STAI) ^39^, which evaluates responses to stressful situations in addition to overall emotional state; and 3) “quality of life” set using the World Health Organization Quality of Life assessment (WHOQOL) ^40^.

**Figure 1.**
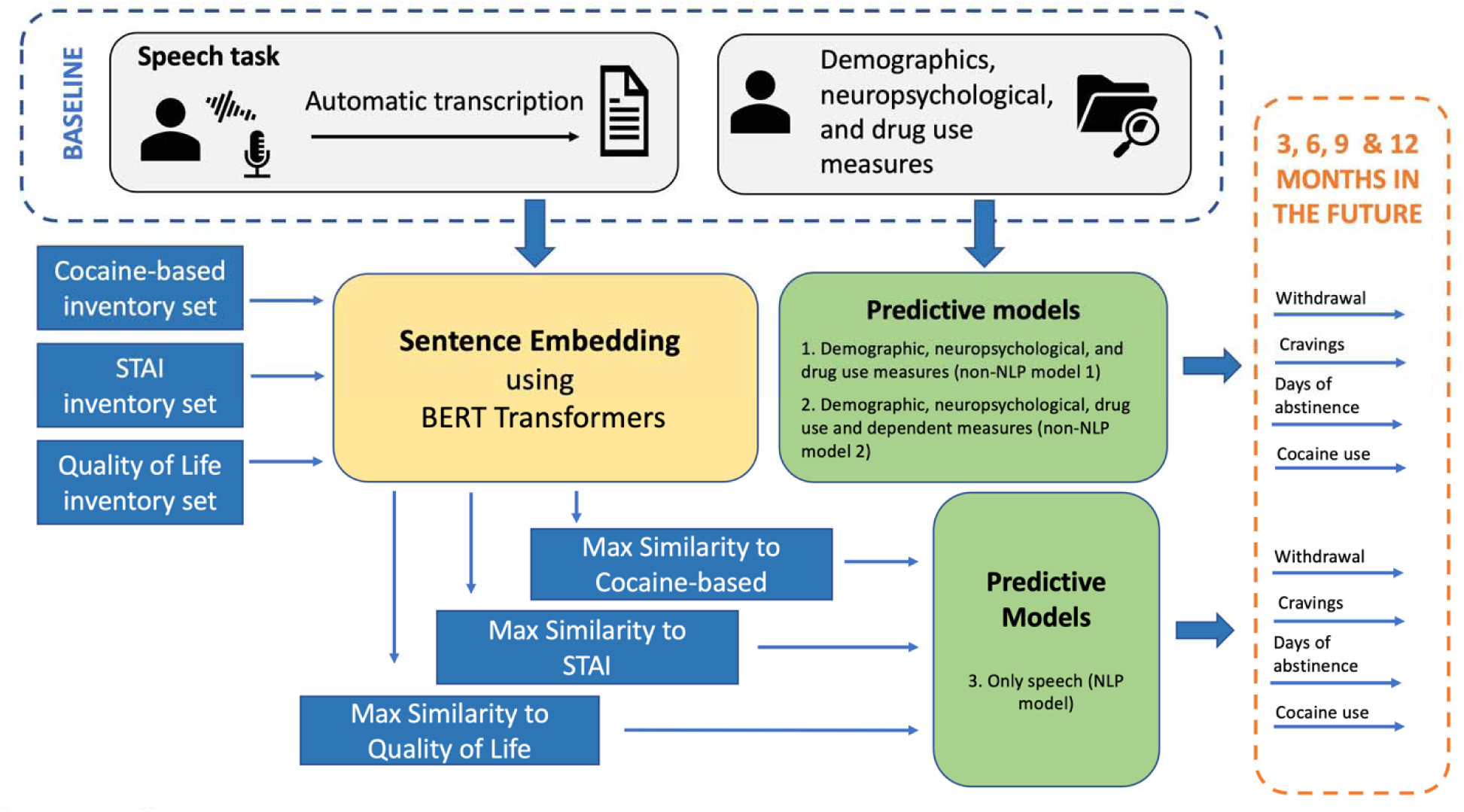
Approach used in this study to analyze recordings of individuals with cocaine use disorders to prospectively predict cocaine-related outcomes at 3, 6, 9, and 12 months follow-up. Under predictive models, model 1 also includes drug use variables listed in the cocaine, alcohol and cigarette use section in Table 1; model 2 in addition includes all dependent measures obtained at baseline (including the four last variables in Table 1). STAI refers to the State-Trait Anxiety Inventory.

Two predictive models did not include NLP data: 1) Non-NLP model 1, which used the baseline demographic, neuropsychological and non-cocaine drug use measures (see Table 1); and 2) Non-NLP model 2, which used all features in Model 1 in addition to the four cocaine use metrics obtained at baseline (see last cluster of variables in Table 1), which also served as our dependent measures (i.e., those obtained at all follow-up sessions). Features derived from the speech variables were entered into NLP model 3. All models were computed independently for the PC and NC prompts.

Since we were interested in predicting future outcomes, only the baseline visit information was used to predict the four dependent measures obtained during each of the subsequent visits at 3 (N=50 participants), 6 (N=36 participants), 9 (N=35 participants), and 12 (N=25 participants) months after baseline across all non-missing subjects.

In all models, feature selection was determined based on rankings in the training set (p>0.1 using a t-test). Given the different scales of the features, two regression algorithms (support vector regression and linear regression) were used to achieve the highest performance (as measured by Spearman correlation coefficients). All generated models were validated using a 10-fold cross-validation. We compared model performance with respect to the null hypothesis (H0: r=0) using a t-test and a Fisher’s test (see supplemental material for further detail). Additionally, to compare our long term results with the previous relevant study ^18^, we binarized the abstinence days outcome measure using 30 days as our threshold (44% participants were labeled abstinent).

## Results

### Model Performance

At the three and six month visits, the non-NLP Model 2 (based on all but the speech features, see Table 1) showed the best performance (r>0.45 and r>0.37, respectively), outperforming the NLP Model 3 for craving prediction at three (*P*=4×10^-4^) and six (*P* =2×10^-2^) months (the highest weighted feature was craving at baseline) with a similar result for abstinence length at the three months’ time point (*P* =2×10^-2^; the predictive feature was abstinence length at baseline). At nine months, all the models performed similarly except for the non-NLP Model 2, which better predicted days of abstinence but not statistically significant with respect to the other models (Fisher’s test *P* =1×10^-1^). At 12 months and across the PC and NC prompts, the NLP Model 3 predicted all four dependent variables: withdrawal (r=0.43, *P* =3×10^-2^), craving (r=0.72, *P* =5×10^-5^), days of abstinence (r=0.76, *P* =1×10^-5^), and cocaine use in the past 90 days (r=0.61, *P* =2×10^-3^), accounting for almost 50% of the variance in drug use behaviors one year later (for craving and abstinence days) (Figure 2), performing better than the other models (by Fisher’s test) for abstinence length. For predicting this measure (based on its binarization at 12 months to > or =<30 days) and using NLP model 3, we achieved AUC of 0.92 (Accuracy = 0.84). Of note, the best predictive models for symptoms of withdrawal and craving were based on the PC prompt, whereas the best predictive models for recent drug use patterns (days of abstinence and drug use in the previous 90 days) were based on the NC prompt. Additional results using models focusing on the cocaine-related inventory set or combination of NC and PC features are described in the supplement.

**Figure 2:**
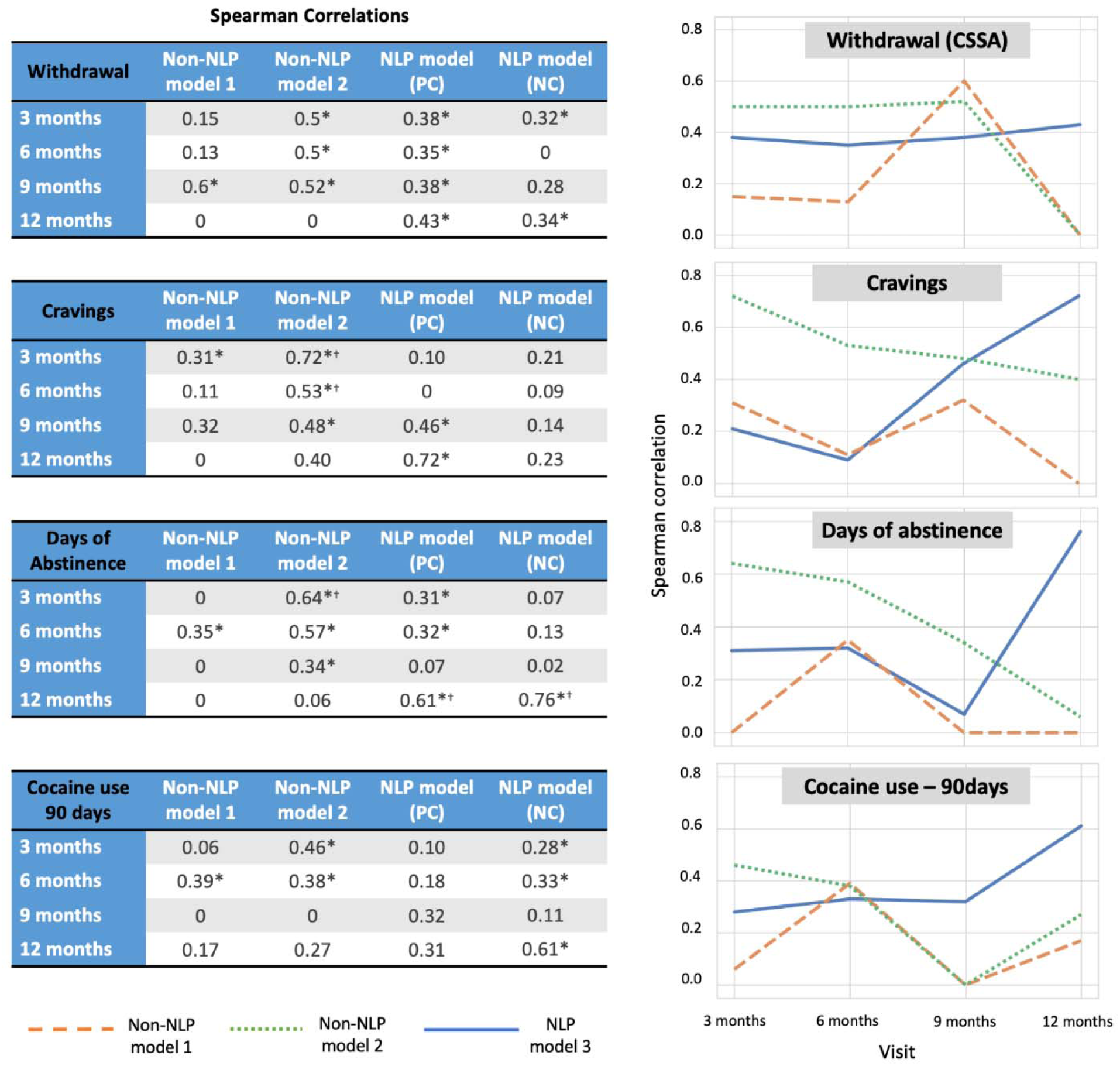
Prediction of longitudinal outcomes in individuals with cocaine use disorder. Tables at the left show the performance of the models that include baseline demographic, neuropsychological and drug use measures (non-NLP Model 1), in addition to the dependent/to be predicted measures obtained at baseline (non-NLP Model 2) and NLP models for positive consequences of abstinence (PC) and negative consequences of drug use (NC) for predicting four selected drug use measures (withdrawal, craving, abstinence days, and cocaine use in the last 90 days) at subsequent visits: 3 (N=50), 6 (N=36), 9 (N=35) and 12 (N=25) months after baseline. Symbols next to the values indicate that the model is statistically significant with regard to the null hypothesis (*) and when comparing between non-NLP and NLP models (^†^). Plots on the right show a different representation of the tables (non-NLP Model 1 = an orange dashed line; non-NLP Model 2= a green dotted line; NLP models (the best of the NC or PC prompts) = a blue solid line).

### Feature Weights

Here we report the weights assigned by the best performing NLP-based models at 12 months (linear regression models showed better performance than the support vector regression models except for withdrawal). The highest weights for withdrawal (from the PC speech) were dominated by the similarity to “I am happy” (predicting less withdrawal) and “I do not love myself” (predicting more withdrawal) (Figure 3a). Weights for craving (also from the PC speech) were dominated by “I do not love myself” and other items related to self-image (“I have support from my friends” and “I want a better life”), all predicting more craving (Figure 3b). For days of abstinence, the highest weights (from the NC speech) were for “I am tense”, predicting fewer days of abstinence; conversely, similarity to “I am not able to concentrate” predicted more days of abstinence (Figure 4a). Finally, the only significant weight for predicting more frequent use in the past 90 days was similarity to “I have enough money to meet my needs” (from the NC speech) (Figure 4b).

**Figure 3:**
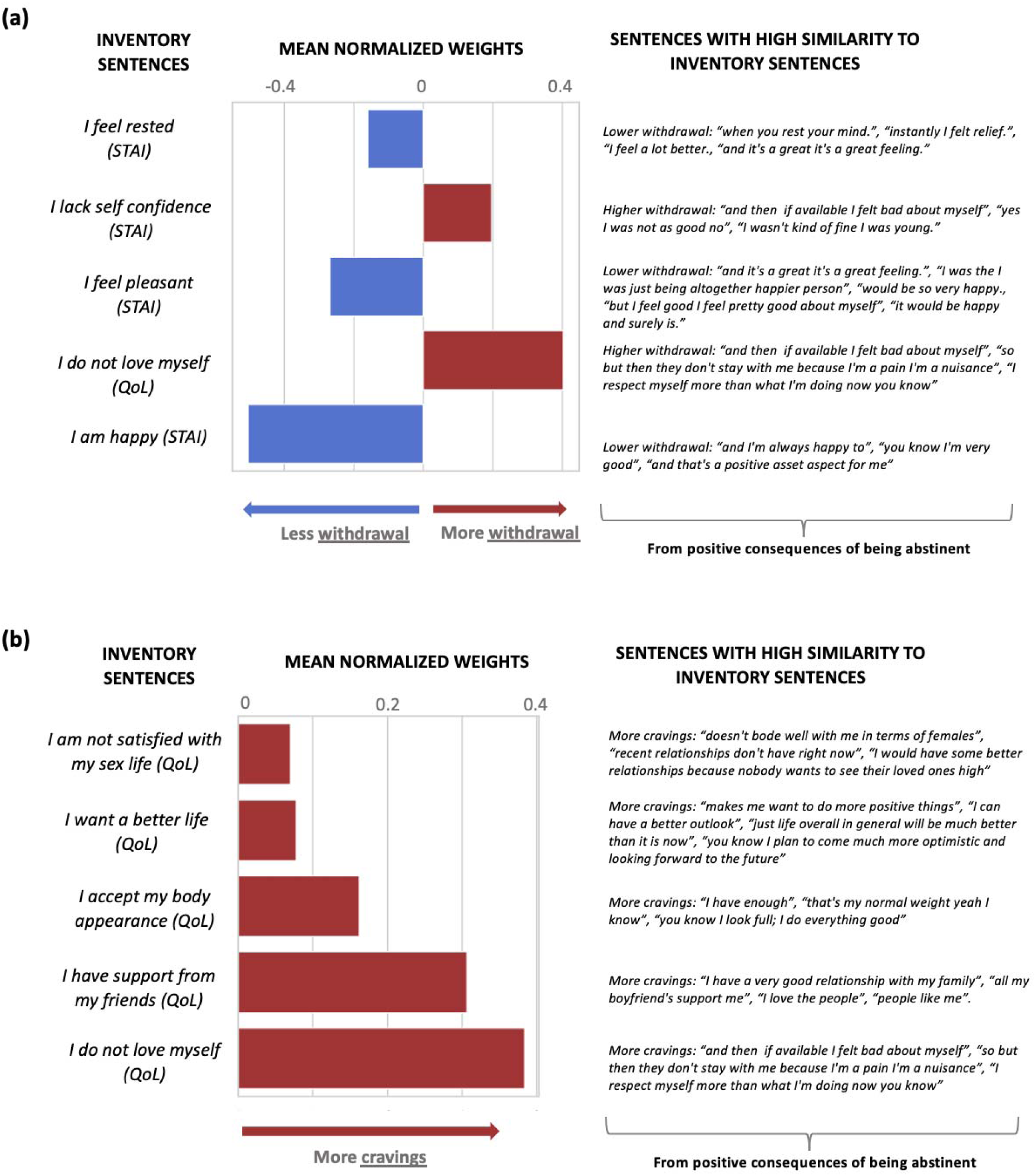
Predictive features of the first visit interview for: (a) withdrawal in the fifth visit (12 months), (b) craving in the fifth visit (12 months). Larger positive weights predict <u>more</u> withdrawal symptoms and craving. Both models are based on the PC section of the interviews.

**Figure 4:**
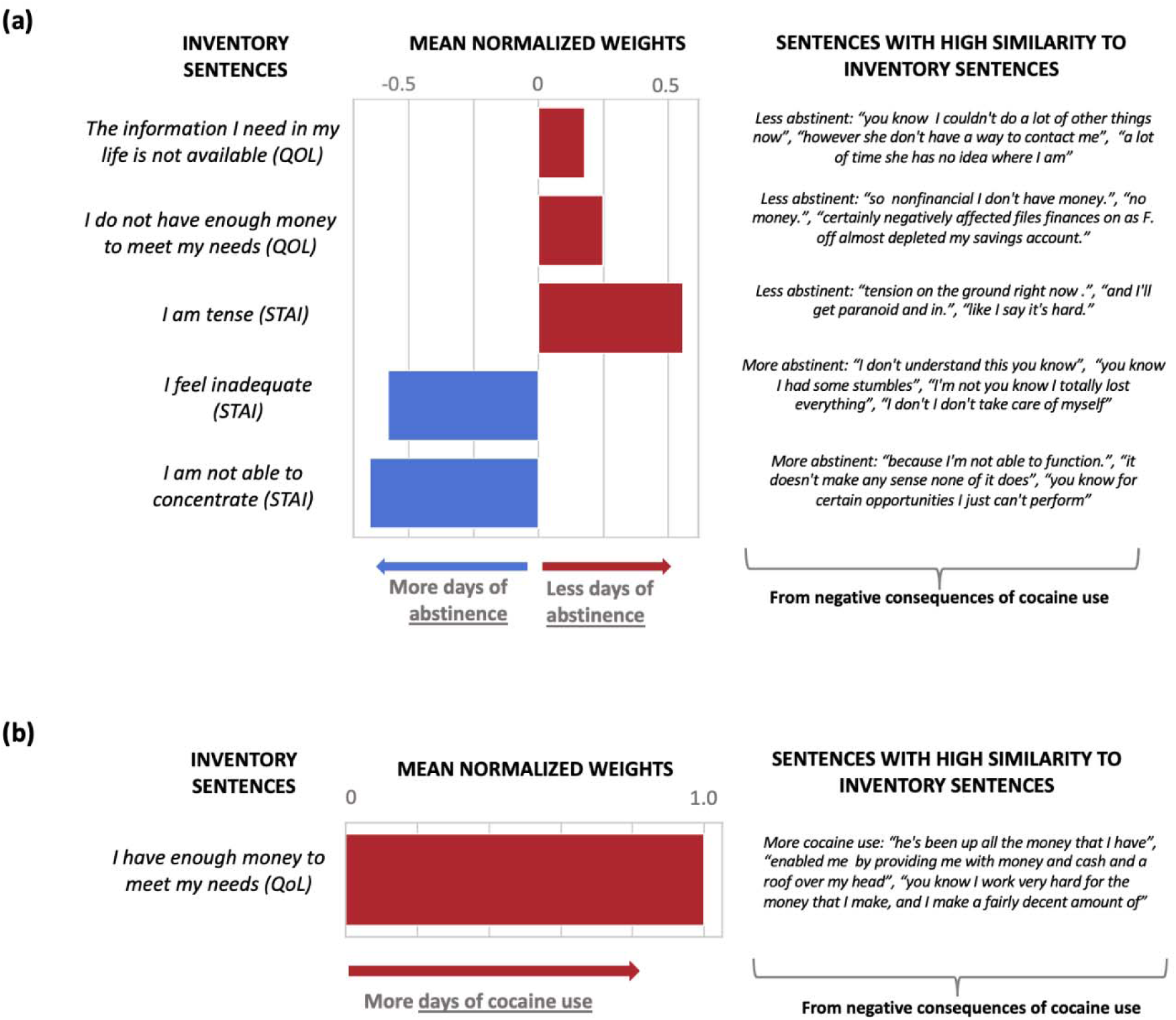
Predictive features of the first visit interview for: (a) days of abstinence in the fifth visit (12 months), (b) the previous 90 days of cocaine use in the fifth visit (12 months). Larger positive weights predict days of abstinence and more days of cocaine use. Both models are based on the NC section of the interviews.

## Discussion

In this longitudinal study, we explored the potential of baseline linguistic markers, as compared to common demographic, neuropsychological and drug use measures, to prospectively predict drug use and addiction symptomatology at three-month intervals for up to one year in initially abstinent iCUD. Our results demonstrated differential predictive outcomes for the NLP-based models as directly compared to non-NLP models as a function of time. Specifically, for predicting outcomes at three and six-months timepoints, the comprehensive non-NLP Model 2, consisting of all measures collected at baseline (including the to be predicted drug use measures), outperformed the NLP Model (and the non-NLP Model 1, consisting of only the demographic and neuropsychological measures), suggesting a degree of continuity over time of drug use patterns and symptoms (especially of craving and abstinence length) with respect to their values at the baseline visit. Importantly, the NLP Model, consisting of speech about personal experiences with cocaine use and abstinence, outperformed the non-NLP models at predicting the 12 months outcomes. Symptoms of withdrawal and craving at 12 months were best predicted by the PC speech, whereas abstinence duration and cocaine use frequency in the previous 90 days were best predicted by the NC speech. For craving and abstinence, these NLP models predicted almost half of the variance one year after baseline, achieving classification rates of 84% for the latter.

Cognitive and behavioral impairments, including in inhibitory control/impulsivity, salience attribution, memory, and decision making, have been consistently and reliably documented in individuals with SUD^41–43^. A growing body of evidence suggests that, when compared to standard clinical interviews and other tools, effective measures of these domains, using validated behavioral tasks and neuroimaging, can provide highly sensitive predictors of future drug use patterns in people with SUD ^44^. In search of such predictors, laboratory-based well-controlled tasks are usually chosen as they are perceived to offer precise and specific insights into drug addiction and recovery mechanisms. However, given these tasks’ limitations (e.g., in sampling from representative and large populations, questionable replicability and generalizability to naturalistic behaviors), here we chose speech and NLP techniques that can quantify complex contextual features of language not immediately apparent to human observers ^45^. For the first time, we demonstrate that, used with sophisticated analysis tools, language can predict longitudinal drug use outcomes in people with SUD initially abstaining from cocaine. Given the ubiquity of language, our NLP results represent an important step towards fine-tuning the use of language and spontaneous speech in both basic and clinical studies, with applications to developing targeted prevention and treatment efforts, in drug addiction.

Our results suggest that linguistic features can better predict long-term outcomes, whereas other, non-NLP, variables may be better suited for shorter-term predictions. The predictive value of these non-NLP variables at shorter timepoints may derive from their stable and/or slowly evolving dynamic of change, best captured over limited durations (i.e., drug use patterns encompassing abstinence and symptoms such as craving are likely to be more similar at three-month intervals than after 12 months). In contrast, consistent with the representation of language in higher-order/more abstract thought processes such as cognitive reappraisal ^46^ that develop during prolonged abstinence and recovery ^47^, features of speech at baseline may capture individual variability that can better predict outcomes at longer intervals. It remains to be established whether, and how, this observation maps onto the nonlinear patterns of change with abstinence in certain cognitive/motivational and behavioral processes encompassing the incubation of craving effect, whereby following a period of withdrawal, cue-induced drug seeking progressively increases before beginning to decline ^48–51^; a similar effect has been suggested for inhibitory control^52^ as potentially generalizable to other higher-order cognitive functions. A study in inpatients with SUD, where abstinence is maintained as ascertained longitudinally, is needed to answer these questions.

Interestingly, the most significant features for the 12-month prediction were derived from speech similarity to the quality-of-life assessments, and not to the cocaine-related inventories. This observation supports the use of the abstract embeddings in NLP analyses, further encouraging reliance on non-direct drug use measures as primary predictors and/or dependent measures in clinical trials in SUD ^5^. Our findings also suggest differential effects of language modeling on outcome prediction depending on the context (positive or negative) of speech. In our study, speech that related the positive effects of abstaining from drugs (PC speech) predicted craving and withdrawal. Specifically, similarity to statements such as “I am happy” was associated with less withdrawal, whereas similarity to statements such as “I do not love myself” was associated with more withdrawal. For craving, some of the results were unexpected (e.g., similarity to the latter statement was the most predictive of higher craving scores, which were also predicted by similarity to statements such as “I have support from my friends” and “I want a better life”), as remains to be further explored vis-à-vis impairments in self-awareness/insight into the severity of one’s illness ^53, 54^. Speech that related the negative consequences of using drugs (NC speech) predicted abstinence duration and cocaine use in the past 90 days. Specifically, similarity to statements such as “I am tense” or “not able to concentrate” was associated with shorter and longer abstinence duration, respectively, suggesting a relationship between poor emotional/cognitive function with recent drug use.

Finally, similarity to the statement “I have enough money to meet my needs” was associated with more frequent use in the past 90 days, which could reflect economics of SUD in the wild (e.g., greater access to drugs when one’s financial situation is stable). Overall, our results suggest specificity of the context of these statements in predicting disparate drug use outcomes, implying the use of positive self-talk (e.g., envisioning one’s future in a more positive way or enhancing one’s self-image and/or perceived cognitive function) in reductions in withdrawal symptoms (and potentially also reduced craving and longer abstinence) 12 months later.

There are a few limitations of this study that are important to address. First, the relatively small size of our sample could have precluded the emergence of important predictive features, clearly necessitating replication in a larger sample. In a related vein, subject attrition at each successive time point may have limited our ability to observe effects at the longer time intervals and may have inflated our classification accuracy rates. Second, the NLP analysis was limited to data collected at a single baseline time point; changes in speech during abstinence may reveal further insights into the recovery process, which warrant an in-depth independent investigation; the contribution of the acoustic features to results also remains to be explored. Third, it is important to consider that speech relating to one’s lived experiences may be especially sensitive to demographic factors such as age, race, sex, education, and socioeconomic status, factors that could also modulate relapse susceptibility (e.g., it is enhanced by the male gender and not participating in follow-up SUD treatment ^55^). Our sample consisted of >80% men and African American individuals mostly between 40 and 60 years of age living in the New York City metropolitan area, potentially reducing sensitivity (encompassing the limited performance of the non-NLP Model 1), and limiting generalizability of our results to different/more diverse populations.

One of the biggest impediments for precisely modeling human addiction recovery across time is the lack of dense and longitudinal measurements of individuals with SUD, especially during the first year from an abstinent baseline, the time of most drastic changes in drug use. A complex brain function, speech conveys people’s emotions, thoughts, and perceptions. It is an ecologically valid outcome measure people use daily, for the duration of the day. Indeed, speech is as natural as some of the most basic of human bodily functions. We report that NLP of minimally-constrained short and spontaneous speech samples obtained at an initially abstaining baseline from iCUD can be used to predict repeatedly sampled outcomes, encompassing withdrawal, craving, days of abstinence and cocaine use, with best predictions for outcomes assessed a full year later. The ability to predict outcomes is essential for the ultimate tailoring of addiction treatment programs to individuals’ specific needs, to enhance relapse prevention strategies and improve the overall effectiveness of substance abuse interventions. By leveraging language-based predictive models of contextualized speech, that could be obtained at scale (e.g., with mobile devices at rehabilitation centers), healthcare providers could optimize care, reduce the risk of relapse, and support individuals on their own path to recovery.

## Acknowledgement

We are very grateful to the individuals who participated in this longitudinal study and whose responses were key for this work. We also thank John Gray for his help with the database and Ahmet Ceceli for his insights on the paper.

## Author Contributions

The corresponding author, Rita Z. Goldstein, had full access to all the data in the study and take responsibility for the integrity of the data and the accuracy of the data analysis.

*Concept and design:* Carla Agurto, Guillermo Cecchi, Rita Z. Goldstein

*Acquisition, analysis, or interpretation of data:* Carla Agurto, Guillermo Cecchi, Sarah King, Rita Z. Goldstein.

*Drafting of the manuscript:* Carla Agurto, Guillermo Cecchi, Sarah King, Rita Z. Goldstein

*Critical revision of the manuscript for important intellectual content:* Carla Agurto, Guillermo Cecchi, Sarah King, Elif K. Eyigoz, Muhammad A. Parvaz, Nelly Alia-Klein, Rita Z. Goldstein.

*Statistical analysis:* Carla Agurto, Guillermo Cecchi, Rita Z. Goldstein

*Administrative, technical, or material support:* Carla Agurto, Guillermo Cecchi, Sarah King, Elif K. Eyigoz, Muhammad A. Parvaz, Nelly Alia-Klein, Rita Z. Goldstein.

*Supervision:* Rita Z. Goldstein

## References

1. Yip SW, Konova AB. Densely sampled neuroimaging for maximizing clinical insight in psychiatric and addiction disorders. Neuropsychopharmacology. 2022;47(1):395–396. doi:10.1038/s41386-021-01124-0

2. Huang Y, Ceceli AO, Kronberg G, et al. Cortico-striatal engagement during cue-reactivity, reappraisal, and savoring of drug and non-drug stimuli predicts craving in heroin addiction. Published online 2023.

3. Parvaz MA, Malaker P, Zilverstand A, Moeller SJ, Alia-Klein N, Goldstein RZ. Attention bias modification in drug addiction: Enhancing control of subsequent habits. Proceedings of the National Academy of Sciences. 2021;118(23):e2012941118. doi:10.1073/pnas.2012941118

4. Goldstein RZ, Tomasi D, Alia-Klein N, et al. Dopaminergic Response to Drug Words in Cocaine Addiction. J Neurosci. 2009;29(18):6001–6006. doi:10.1523/JNEUROSCI.4247-08.2009

5. Goldstein RZ. Neuropsychoimaging Measures as Alternatives to Drug Use Outcomes in Clinical Trials for Addiction. JAMA Psychiatry. 2022;79(9):843–844. doi:10.1001/jamapsychiatry.2022.1970

6. Afshar M, Sharma B, Dligach D, et al. Development and multimodal validation of a substance misuse algorithm for referral to treatment using artificial intelligence (SMART-AI): a retrospective deep learning study. The Lancet Digital Health. 2022;4(6):e426–e435. doi:10.1016/S2589-7500(22)00041-3

7. Blackley SV, MacPhaul E, Martin B, Song W, Suzuki J, Zhou L. Using Natural Language Processing and Machine Learning to Identify Hospitalized Patients with Opioid Use Disorder. AMIA Annu Symp Proc. 2021;2020:233–242.

8. Hastings JS, Howison M, Inman SE. Predicting high-risk opioid prescriptions before they are given. Proc Natl Acad Sci U S A. 2020;117(4):1917–1923. doi:10.1073/pnas.1905355117

9. Shah-Mohammadi F, Cui W, Bachi K, Hurd Y, Finkelstein J. Using Natural Language Processing of Clinical Notes to Predict Outcomes of Opioid Treatment Program. Annu Int Conf IEEE Eng Med Biol Soc. 2022;2022:4415–4420. doi:10.1109/EMBC48229.2022.9871960

10. Ni Y, Bachtel A, Nause K, Beal S. Automated detection of substance use information from electronic health records for a pediatric population. Journal of the American Medical Informatics Association. 2021;28(10):2116–2127. doi:10.1093/jamia/ocab116

11. Goodman-Meza D, Tang A, Aryanfar B, et al. Natural Language Processing and Machine Learning to Identify People Who Inject Drugs in Electronic Health Records. Open Forum Infect Dis. 2022;9(9):ofac471. doi:10.1093/ofid/ofac471

12. Goodwin TR, Maldonado R, Harabagiu SM. Automatic recognition of symptom severity from psychiatric evaluation records. J Biomed Inform. 2017;75S:S71–S84. doi:10.1016/j.jbi.2017.05.020

13. Joyce C, Markossian TW, Nikolaides J, et al. The Evaluation of a Clinical Decision Support Tool Using Natural Language Processing to Screen Hospitalized Adults for Unhealthy Substance Use: Protocol for a Quasi-Experimental Design. JMIR Res Protoc. 2022;11(12):e42971. doi:10.2196/42971

14. Ridgway JP, Uvin A, Schmitt J, et al. Natural Language Processing of Clinical Notes to Identify Mental Illness and Substance Use Among People Living with HIV: Retrospective Cohort Study. JMIR Med Inform. 2021;9(3):e23456. doi:10.2196/23456

15. Goodman-Meza D, Shover CL, Medina JA, Tang AB, Shoptaw S, Bui AAT. Development and Validation of Machine Models Using Natural Language Processing to Classify Substances Involved in Overdose Deaths. JAMA Network Open. 2022;5(8):e2225593. doi:10.1001/jamanetworkopen.2022.25593

16. Ding T, Hasan F, Bickel WK, Pan S. Interpreting Social Media-Based Substance Use Prediction Models with Knowledge Distillation. In: 2018 IEEE 30th International Conference on Tools with Artificial Intelligence (ICTAI).; 2018:623–630. doi:10.1109/ICTAI.2018.00100

17. Yang G, King, Sarah G, Lin, Hung-Mo, Goldstein, Rita Z. Emotional expression on social media support forums for substance cessation: Observational study of text-based Reddit posts. JMIR Preprints. Published 2023. Accessed June 26, 2023. https://preprints.jmir.org/preprint/45267

18. Cox DJ, Garcia-Romeu A, Johnson MW. Predicting changes in substance use following psychedelic experiences: natural language processing of psychedelic session narratives. The American Journal of Drug and Alcohol Abuse. 2021;47(4):444–454. doi:10.1080/00952990.2021.1910830

19. Pickens CL, Airavaara M, Theberge F, Fanous S, Hope BT, Shaham Y. Neurobiology of the incubation of drug craving. Trends Neurosci. 2011;34(8):411–420. doi:10.1016/j.tins.2011.06.001

20. Anand T, Kandasamy A, Suman LN. Self-stigma, hope for future, and recovery: An exploratory study of men with early-onset substance use disorder. Ind Psychiatry J. 2022;31(2):299–305. doi:10.4103/ipj.ipj_52_21

21. Liu Y, Wang L, Yu C, et al. How drug cravings affect metacognitive monitoring in methamphetamine abusers. Addictive Behaviors. 2022;132:107341. doi:10.1016/j.addbeh.2022.107341

22. Lauvsnes ADF, Gråwe RW, Langaas M. Predicting Relapse in Substance Use: Prospective Modeling Based on Intensive Longitudinal Data on Mental Health, Cognition, and Craving. Brain Sciences. 2022;12(7):957. doi:10.3390/brainsci12070957

23. Hallgren KA. Remotely Assessing Mechanisms of Behavioral Change in Community Substance Use Disorder Treatment to Facilitate Measurement-Informed Care: Pilot Longitudinal Questionnaire Study. JMIR Form Res. 2022;6(11):e42376. doi:10.2196/42376

24. Steele VR, Fink BC, Maurer JM, et al. Brain Potentials Measured During a Go/NoGo Task Predict Completion of Substance Abuse Treatment. Biol Psychiatry. 2014;76(1):75–83. doi:10.1016/j.biopsych.2013.09.030

25. Steele VR, Maurer JM, Arbabshirani MR, et al. Machine Learning of Functional Magnetic Resonance Imaging Network Connectivity Predicts Substance Abuse Treatment Completion. Biological Psychiatry: Cognitive Neuroscience and Neuroimaging. 2018;3(2):141–149. doi:10.1016/j.bpsc.2017.07.003

26. Luo SX, Martinez D, Carpenter KM, Slifstein M, Nunes EV. Multimodal predictive modeling of individual treatment outcome in cocaine dependence with combined neuroimaging and behavioral predictors. Drug and Alcohol Dependence. 2014;143:29–35. doi:10.1016/j.drugalcdep.2014.04.030

27. Clark VP, Beatty GK, Anderson RE, et al. Reduced fMRI activity predicts relapse in patients recovering from stimulant dependence. Hum Brain Mapp. 2014;35(2):414–428. doi:10.1002/hbm.22184

28. Yip SW, Scheinost D, Potenza MN, Carroll KM. Connectome-Based Prediction of Cocaine Abstinence. AJP. 2019;176(2):156–164. doi:10.1176/appi.ajp.2018.17101147

29. Agurto C, Norel R, Pietrowicz M, et al. Speech markers for clinical assessment of cocaine users. Proc IEEE Int Conf Acoust Speech Signal Process. 2019;2019:6391–6394. doi:10.1109/icassp.2019.8682691

30. Agurto C, Cecchi GA, Norel R, et al. Detection of acute 3,4-methylenedioxymethamphetamine (MDMA) effects across protocols using automated natural language processing. Neuropsychopharmacology. 2020;45(5):823–832. doi:10.1038/s41386-020-0620-4

31. Liu Y, Ott M, Goyal N, et al. RoBERTa: A Robustly Optimized BERT Pretraining Approach. ArXiv. Published online July 26, 2019. Accessed April 11, 2023. https://www.semanticscholar.org/paper/RoBERTa%3A-A-Robustly-Optimized-BERT-Pretraining-Liu-Ott/077f8329a7b6fa3b7c877a57b81eb6c18b5f87de

32. Lokala U, Phukan OC, Dastidar TG, Lamy F, Daniulaityte R, Sheth A. “Can We Detect Substance Use Disorder?”: Knowledge and Time Aware Classification on Social Media from Darkweb. Published online April 20, 2023. Accessed May 16, 2023. http://arxiv.org/abs/2304.10512

33. Kampman KM, Volpicelli JR, McGinnis DE, et al. Reliability and validity of the Cocaine Selective Severity Assessment. Addict Behav. 1998;23(4):449–461. doi:10.1016/s0306-4603(98)00011-2

34. Sobell LC, Sobell M. Timeline Followback Method (Drugs, Cigarettes, and Marijuana). Published 1996. Accessed April 18, 2023. https://www.emcdda.europa.eu/drugs-library/timeline-followback-method-drugs-cigarettes-and-marijuana_en

35. IBM Watson Speech to Text - Overview. Published April 6, 2023. Accessed April 11, 2023. https://www.ibm.com/cloud/watson-speech-to-text

36. Gossop M, Griffiths P, Powis B, Strang J. Severity of dependence and route of administration of heroin, cocaine and amphetamines. Br J Addict. 1992;87(11):1527–1536. doi:10.1111/j.1360-0443.1992.tb02660.x

37. Gossop M, Darke S, Griffiths P, et al. The Severity of Dependence Scale (SDS): psychometric properties of the SDS in English and Australian samples of heroin, cocaine and amphetamine users. Addiction. 1995;90(5):607–614. doi:10.1046/j.1360-0443.1995.9056072.x

38. Sussner BD, Smelson DA, Rodrigues S, Kline A, Losonczy M, Ziedonis D. The validity and reliability of a brief measure of cocaine craving. Drug Alcohol Depend. 2006;83(3):233–237. doi:10.1016/j.drugalcdep.2005.11.022

39. Spielberger CD. State-Trait Anxiety Inventory. In: *The Corsini Encyclopedia of Psychology*. John Wiley & Sons, Ltd; 2010:1–1. doi:10.1002/9780470479216.corpsy0943

40. The World Health Organization Quality of Life (WHOQOL). Accessed April 11, 2023. https://www.who.int/publications-detail-redirect/WHO-HIS-HSI-Rev.2012.03

41. Goldstein RZ, Leskovjan AC, Hoff AL, et al. Severity of neuropsychological impairment in cocaine and alcohol addiction: association with metabolism in the prefrontal cortex. Neuropsychologia. 2004;42(11):1447–1458. doi:10.1016/j.neuropsychologia.2004.04.002

42. Kozak K, Lucatch AM, Lowe DJE, Balodis IM, MacKillop J, George TP. The neurobiology of impulsivity and substance use disorders: implications for treatment. Ann N Y Acad Sci. 2019;1451(1):71–91. doi:10.1111/nyas.13977

43. Verdejo-García A, Lawrence AJ, Clark L. Impulsivity as a vulnerability marker for substance-use disorders: review of findings from high-risk research, problem gamblers and genetic association studies. Neurosci Biobehav Rev. 2008;32(4):777–810. doi:10.1016/j.neubiorev.2007.11.003

44. Yip SW, Kiluk B, Scheinost D. Toward Addiction Prediction: An Overview of Cross-Validated Predictive Modeling Findings and Considerations for Future Neuroimaging Research. Biological Psychiatry: Cognitive Neuroscience and Neuroimaging. 2020;5(8):748–758. doi:10.1016/j.bpsc.2019.11.001

45. Mota NB, Weissheimer J, Finger I, Ribeiro M, Malcorra B, Hübner L. Speech as a graph: developmental perspectives on the organization of spoken language. Biol Psychiatry Cogn Neurosci Neuroimaging. Published online April 19, 2023:S2451–9022(23)00098-8. doi:10.1016/j.bpsc.2023.04.004

46. Otto B, Misra S, Prasad A, McRae K. Functional overlap of top-down emotion regulation and generation: An fMRI study identifying common neural substrates between cognitive reappraisal and cognitively generated emotions. Cogn Affect Behav Neurosci. 2014;14(3):923–938. doi:10.3758/s13415-013-0240-0

47. Garland EL, Roberts-Lewis A, Kelley K, Tronnier C, Hanley A. Cognitive and Affective Mechanisms Linking Trait Mindfulness to Craving Among Individuals in Addiction Recovery. Substance Use & Misuse. Published online March 11, 2014. Accessed April 11, 2023. https://www.tandfonline.com/doi/full/10.3109/10826084.2014.850309

48. Grimm JW, Hope BT, Wise RA, Shaham Y. Neuroadaptation. Incubation of cocaine craving after withdrawal. Nature. 2001;412(6843):141-142. doi:10.1038/35084134

49. Lu L, Grimm JW, Dempsey J, Shaham Y. Cocaine seeking over extended withdrawal periods in rats: different time courses of responding induced by cocaine cues versus cocaine priming over the first 6 months. Psychopharmacology (Berl). 2004;176(1):101–108. doi:10.1007/s00213-004-1860-4

50. Lu L, Grimm JW, Hope BT, Shaham Y. Incubation of cocaine craving after withdrawal: a review of preclinical data. Neuropharmacology. 2004;47 Suppl 1:214–226. doi:10.1016/j.neuropharm.2004.06.027

51. Parvaz MA, Moeller SJ, Goldstein RZ. Incubation of Cue-Induced Craving in Adults Addicted to Cocaine Measured by Electroencephalography. JAMA Psychiatry. 2016;73(11):1127–1134. doi:10.1001/jamapsychiatry.2016.2181

52. Anna Zilverstand, Muhammad Parvaz, Scott Moeller, et al. Whole-Brain Resting-State Connectivity Underlying Impaired Inhibitory Control during Early Versus Longer-Term Abstinence in Cocaine Addiction. Mol Psychiatry. Published online in revision.

53. Moeller SJ, Maloney T, Parvaz MA, et al. Impaired insight in cocaine addiction: laboratory evidence and effects on cocaine-seeking behaviour. Brain. 2010;133(5):1484–1493. doi:10.1093/brain/awq066

54. Orfei MD, Robinson RG, Bria P, Caltagirone C, Spalletta G. Unawareness of illness in neuropsychiatric disorders: phenomenological certainty versus etiopathogenic vagueness. Neuroscientist. 2008;14(2):203–222. doi:10.1177/1073858407309995

55. Waite MR, Heslin K, Cook J, Kim A, Simpson M. Predicting substance use disorder treatment follow-ups and relapse across the continuum of care at a single behavioral health center. Journal of Substance Use and Addiction Treatment. 2023;147:208933. doi:10.1016/j.josat.2022.208933

