## Supplementary material for "Speak and you shall predict: speech at initial cocaine abstinence as a biomarker of long-term drug use behavior"

**Participants**

All participants met criteria for cocaine use disorder based on a comprehensive diagnostic interview at baseline, which included the following: Structured Clinical Interview for Diagnostic and Statistical Manual of Mental Disorders, Fourth Edition, Axis I Disorders (SCID) (First et al., 1996), for clinical diagnostic criteria and the Addiction Severity Index (ASI) ^1^, for lifetime drug and alcohol use and severity, in addition to the tools listed in the main text. We also assessed demographics, neuropsychological measures [reading subtest of the Wide Range Achievement Test for verbal IQ, Matrix Reasoning subtest of the Wechsler Abbreviated Scale of Intelligence ^2^ for nonverbal IQ, and Beck’s Depression Inventory ^3^ for depression symptomatology], severity of dependence on nicotine [Fagerström Test for Nicotine Dependence ^4^] and alcohol use [abbreviated Michigan Alcohol Severity Test^5^], and additional cocaine use measures encompassing route of administration, age of onset, duration of regular use [from the diagnostic interview and the Timeline Followback ^6^], and the Severity of Dependence Scale (SDS) ^7,8^. Note that we also assessed the State-Trait Anxiety Inventory (STAI) ^9^, but these responses were not included as predictors (or outcomes) because we used their contents for the speech embeddings (see main text).

**Speech tasks**

Participants performed a minimally structured speech task where they were instructed to verbally address two prompts, describing the positive consequences (PC) of abstinence and the negative consequences (NC) of using cocaine. Verbatim instructions were “Please talk about the POSITIVE consequences of quitting your drug use on your life. For example, you can talk about how quitting taking drugs has affected your personal (family, health), your professional (job), or social (friend and people around you) life” for PC, and “Please talk about the NEGATIVE consequences of your drug use on your life. For example, you can talk about how taking drugs has affected your personal (family, health), your professional (job), or social (friend and people around you) life” for NC. Participants were asked to speak for 5 minutes for each prompt and were encouraged to continue speaking when necessary. All participants spoke in English.

**Analytical approach**

Given the differences in the contexts and instructions (e.g., regret over the past or current experiences is expected to be represent in NC, and hope and expectations for a better future in the PC prompts), these prompts were analyzed separately. We first manually removed the interviewers’ voice from the recordings (despite minimal presence), resulting in average duration of 279.2 +/- 29.9 seconds for NC and 273.6 +/- 32.1 seconds for PC. Then, the speech samples were transcribed using a customized version of the Speech to text IBM API. We performed language and acoustic model customizations for each speaker to improve the accuracy of the transcriptions. More specifically, the language model for transcribing the recordings was developed using an iterative method as follows: we first created a language model using the transcriptions created by the non-customized baseline model of the API. This corpus of 88173 words (from 481 recordings) was in turn used to train a language model to perform customized speech recognition.

These sentences were then embedded into a 1024-dimensional space using BERT encoding, specifically RoBERTa ^10^, for further processing. After performing the sentence embeddings, each response to both prompts was represented as *n*-dimensional vectors whose components were the cosine similarity of its sentences to each inventory item sentence; the cosine similarity is a measure of how similar two sentences are based on the proximity of their embeddings. For our analysis, from each subject we only considered as features the maximum cosine similarity to each inventory sentence. In the final step, these maximum similarity scores were used as input into the regression models.

**Model Performance (additional results)**

Models developed using features derived from only similarity scores with the cocaine-related set of inventories were not superior in any instance to the ones developed with the other inventories. In fact, for predictions at 12 months, the best model using cocaine-related inventories achieved r = 0.21 for days of abstinence. Additionally, we explored combining features for both prompts (PC and NC), performing this analysis for predicting outcomes at 12 months. A higher correlation (r=0.84) was observed for abstinence length (as compared to using single prompts: NC: r=0.76, PC: r=0.61) although this difference did not reach statistical significance (Fisher’s test *P*=2x10^-1^). An additional feature (with a negative weight) in this new model was “I lack self confidence”, which predicted greater withdrawal.

**References**

1.	McLellan AT, Kushner H, Metzger D, et al. The Fifth Edition of the Addiction Severity Index. *J Subst Abuse Treat*. 1992;9(3):199-213. doi:10.1016/0740-5472(92)90062-s

2.	Wechsler D. *Wechsler Abbreviated Scale of Intelligence WASI: Manual*. Pearson/PsychCorpl; 1999. https://books.google.com/books?id=adTXtwAACAAJ

3.	Beck AT, Steer RA, Carbin MG. Psychometric properties of the Beck Depression Inventory: Twenty-five years of evaluation. *Clinical Psychology Review*. 1988;8(1):77-100. doi:10.1016/0272-7358(88)90050-5

4.	Heatherton TF, Kozlowski LT, Frecker RC, Fagerström KO. The Fagerström Test for Nicotine Dependence: a revision of the Fagerström Tolerance Questionnaire. *Br J Addict*. 1991;86(9):1119-1127. doi:10.1111/j.1360-0443.1991.tb01879.x

5.	Selzer ML. The Michigan Alcoholism Screening Test: The Quest for a New Diagnostic Instrument. *AJP*. 1971;127(12):1653-1658. doi:10.1176/ajp.127.12.1653

6.	Sobell LC, Sobell M. Timeline Followback Method (Drugs, Cigarettes, and Marijuana). Published 1996. Accessed April 18, 2023. https://www.emcdda.europa.eu/drugs-library/timeline-followback-method-drugs-cigarettes-and-marijuana_en

7.	Gossop M, Griffiths P, Powis B, Strang J. Severity of dependence and route of administration of heroin, cocaine and amphetamines. *Br J Addict*. 1992;87(11):1527-1536. doi:10.1111/j.1360-0443.1992.tb02660.x

8.	Gossop M, Darke S, Griffiths P, et al. The Severity of Dependence Scale (SDS): psychometric properties of the SDS in English and Australian samples of heroin, cocaine and amphetamine users. *Addiction*. 1995;90(5):607-614. doi:10.1046/j.1360-0443.1995.9056072.x

9.	Spielberger CD. State-Trait Anxiety Inventory. In: *The Corsini Encyclopedia of Psychology*. John Wiley & Sons, Ltd; 2010:1-1. doi:10.1002/9780470479216.corpsy0943

10.	Liu Y, Ott M, Goyal N, et al. RoBERTa: A Robustly Optimized BERT Pretraining Approach. *ArXiv*. Published online July 26, 2019. Accessed April 11, 2023. https://www.semanticscholar.org/paper/RoBERTa%3A-A-Robustly-Optimized-BERT-Pretraining-Liu-Ott/077f8329a7b6fa3b7c877a57b81eb6c18b5f87de
